# Small Cell Lung Cancer Neuroendocrine Subtypes are Associated with Different Immune Microenvironment and Checkpoint Molecule Distribution

**DOI:** 10.1101/2020.02.02.930305

**Authors:** David Dora, Christopher Rivard, Hui Yu, Paul Bunn, Kenichi Suda, Shengxiang Ren, Shivaun Lueke Pickard, Viktoria Laszlo, Tunde Harko, Zsolt Megyesfalvi, Judit Moldvay, Fred R. Hirsch, Balazs Dome, Zoltan Lohinai

## Abstract

Small cell lung cancer (SCLC) has recently been sub-categorized into neuroendocrine (NE)- high and NE-low subtypes showing ‘immune desert’ and ‘immune oasis’ phenotypes, respectively. We aimed to characterize the immune cell localization and the microenvironment according to immune checkpoints and NE subtypes in human SCLC tissue samples at the protein level. In this cross-sectional study, we included 32 primary tumors and matched lymph node (LN) metastases of resected early-stage, histologically confirmed SCLC patients, which were previously clustered into NE subtypes using NE-associated key RNA genes. Immunohistochemistry (IHC) was performed on FFPE TMAs with antibodies against CD45, CD3, CD8 and immune checkpoints including poliovirus receptor (PVR) and Indoleamine 2,3-dioxygenase (IDO).

According to our results, the stroma was significantly more infiltrated by immune cells both in primary tumors and LN metastases (vs tumor cell nests). Immune (CD45+) cell density was significantly higher in tumor nests (110.6 ± 24.95 vs 42.74 ± 10.30, cell/mm2, p= 0.0048), with increased CD8+ effector T cell infiltration (21.81 ± 5.458 vs 3.16 ± 1.36 cell/mm2, p < 0.001) in NE-low vs NE-high tumors. Furthermore, the expression of IDO was confirmed on stromal and endothelial cells, and it positively correlated (r= 0.755, p<0.01) with higher immune cell density both in primary tumors and LN metastases, regardless of the NE pattern. Expression of IDO in tumor nests was significantly higher in NE-low (vs NE-high) primary tumors. PVR expression was significantly higher in NE-low (vs NE-high) patients both in primary tumors) and LN metastases.

To our knowledge, this is the first human study that demonstrates in situ that NE-low tumors are associated with increased immune cell infiltration compared to NE-high tumors. PVR and IDO are potential new targets in SCLC, with increased expression in the NE-low subtype, providing key insight for further prospective studies on potential biomarkers and targets for SCLC immunotherapies.

## INTRODUCTION

Very recently, there have been initial milestones achieved in understanding small cell lung cancer (SCLC) biology. Two recent randomized trials comparing etoposide-platinum doublet therapy alone to the same therapy plus a checkpoint inhibitor (atezolizumab or durvalumab) as first-line therapy showed significant increases in PFS (4.3-5.2 month), response rate and overall survival (12.3 - 13 vs 10.3 months) with the immunotherapy (Horn et al., 2018; Paz-Ares et al., 2019). However, these benefits are limited, and biomarkers, such as smoking status, tumor mutation burden (TMB), and programmed cell death-ligand 1 (PD-L1) expression, did not predict outcome. The lack of a biomarker and the benefit in a small portion of patients points toward the idea that SCLC might be associated with a different immunological microenvironment (Antonia et al., 2016; Hellmann et al, 2018). Furthermore, a lack of tumor tissue availability due to disease aggressiveness limits our understanding of crucial immunological mechanisms, including immune cell infiltration, inter-tumor, and intra-tumor heterogeneity, which may be one reason behind the long-term failure of immunotherapies. Moreover, in many patients, LN metastases are the primary motivators for rapid disease progression, and their immunological environment is far less understood.

SCLC is no longer considered as a single-disease entity, and subtypes are defined by distinct RNA gene expression profiles and can be classified into neuroendocrine (NE)-high and NE - low tumors, which might have different immunogenicity (Rudin et al, 2019). NE-high is characterized by decreased immune cell infiltration defined as a cold or ‘immune desert’ phenotype, based on low levels of immune cell-related RNA expression. In contrast, NE-low was associated with tumors with increased immunogenicity, in other words ‘hot’ or ‘immune oasis’ phenotype (Calbo et al., 2011; George et al., 2015; McColl et al., 2017; Rudin et al., 2019). Consequently, NE-low SCLC patients may more likely respond to immunotherapies (Gazdar, 2018; Saito et al., 2018). The immune infiltrate is comprised of innate and adaptive immune cells, whose populations are heterogeneous across tumor types and patients, and include non-specific immune cell types, such as macrophages, neutrophil granulocytes, dendritic, mast and natural killer (NK) cells, or effector cells of specific immunity, like Band CD3+ T cells (CD4+ T helper, CD8+ cytotoxic T, and regulatory T [Treg] cells), localized in tumor nests, or adjacent tumor stroma (Fridman et al, 2012). A high number of DCs, NK cells, B cells, and CD8+ T cells was associated with improved prognosis, while the presence of Treg cells correlates with decreased survival time in NSCLC (Soo et al, 2008; Hiraoka et al, 2006). The invasion of tumor nests by immune cells confers better overall survival (OS) in lung cancer and other malignancies (Welsh et al, 2005; Miksch et al, 2019).

In addition to the presence of tumor-infiltrating immune cells, the expression of specific immune checkpoints is also a crucial immune-suppressing factor in many cancers. Poliovirus receptor (PVR), an important factor in the SCLC microenvironment, is an adhesion molecule involved in cell motility, as well as NK cell and T-cell-mediated immunity. PVR is relatively absent in normal tissues, but regularly overexpressed in malignancies promoting tumor cell invasion and migration (Sloan et al, 2004). PVR expression was detected at low levels in multiple cell types of epithelial origin, and overexpressed in cancers of epithelial and neural origins (Yu et al., 2009; Johnston et al., 2014; Chauvin et al., 2015). PVR was also proved to play a crucial role in oncoimmunity, as a ligand of coinhibitory receptor TIGIT and CD96 on NK and T cells (Gao et al, 2017). Recently, it was reported that PVR is highly expressed in SCLC cell lines with minimal expression observed on immune cells in the tumor microenvironment (Hu et al., 2018).

Indoleamine-2,3-dioxygenase 1 (IDO), is a key factor in defining cancer immunogenicity (Hornyák et al, 2018), and is a cytosolic enzyme catalyzing the first and rate-limiting step of tryptophan (Trp) catabolism. Multiple studies revealed that the accumulation of Trp metabolites promotes the differentiation of Treg cells and induces the apoptosis of effector T cells with consequent immunosuppression (Munn et al, 2013; van Baren and Van den Eynde 2015). IDO is overexpressed in many tumor types exploiting immunosuppressive mechanisms to promote their spread and survival (Katz et al, 2008).

This study focuses on the evaluation and quantification of immune cell infiltration by localization and distribution patterns in the stroma and tumor nests according to SCLC NE subtypes. In addition, SCLC tumors are evaluated for emerging immune checkpoints including PVR and IDO to allow for new trials of immune therapy in these SCLC subsets.

## MATERIALS AND METHODS

### Ethics Statement

Research was conducted in accordance with the guidelines of the Helsinki Declaration of the World Medical Association. The approval of the Hungarian Scientific and Research Ethics Committee of the Medical Research Council, (ETTTUKEB-7214-1/2016/EKU) was obtained and waived the need for individual informed consent for this study. After the collection of clinical data, patient identifiers were removed so that patients may not be identified either directly or indirectly.

### Study population

A total of 32 histologically confirmed early-stage SCLC patients with available primary tumor tissue and matched LN metastases were included in our study as previously described (Lohinai et al, 2020). All patients underwent surgical resection in the period from 1978 to 2013 at the National Koranyi Institute of Pulmonology. Formalin-fixed, paraffin-embedded (FFPE) tissue samples from primary tumors and LN metastases were obtained at the time of lung resection surgery. Clinicopathological characteristics were described earlier (Lohinai et al, 2020).

### Tissue processing

SCLC patient tumors were obtained by surgical resection and were fixed and processed into paraffin blocks. TMA construction from FFPE blocks was performed as previously described (Battifora 1986). Briefly, 4-micron sections from each tissue block were prepared using a HM-315 microtome (Microm) and placed on charged glass slides (Colorfrost Plus, #22-230-890, Fisher). Slides were stained for H&E on an automated Tissue-Tek Prisma staining platform (Sukura). H&E slides were reviewed by a laboratory pathologist for tumor area and the tumor boarder marked. Marked-stained sections were used to guide the technician as to the location for punch tissue removal. Two 1-mm punches of tissue were taken from each donor tissue block for primary tumors, and one 1-mm punch from LN metastases blocks and seated into a recipient paraffin block in a positionally-encoded array format (MP10 1.0 mm tissue punch on a manual TMA instrument, Beecher Instruments).

### Molecular analysis

RNA expression data from primary and LN FFPE tumor tissue samples was obtained using the HTG EdgeSeq Targeted Oncology Biomarker Panel as previously described (Lohinai et al., 2020). Tumors were clustered into NE-low (n=21) and NE-high (n=43) subtypes according to their neuroendocrine gene expression patterns as previously reported (Lohinai et al., 2020).

### Immunohistochemistry

Four-micron-thick sections were cut from FFPE TMA blocks for IHC staining. Slides were stained on a Leica Bond RX autostainer using rabbit monoclonal antibody for IDO (#86630), CD45 (#13917), CD3□ (#85061), CD8α (#8112) and PVR/CD155 (#81254). All antibodies were from Cell Signaling and diluted 1:200 with Cell Signaling antibody diluent (#8112) prior to staining. Slides were stained using the Bond Polymer Refine Detection kit (#DS9800) with Leica IHC Protocol F and exposed to epitope retrieval 1 (low pH) for twenty minutes. Following staining, slides were cleared and dehydrated on an automated Tissue-Tek Prisma platform and cover slipped using a Tissue-Tek Film cover slipper. The detection of protein expression was optimized in human tonsil and thymus tissue as a positive control.

### Cell counting and morphometry

Images of TMA sections were captured via a BX53 upright Olympus microscope and a DP74 color CMOS camera with 10× magnification objectives in 20MP resolution for scoring and cell counting, and with 20× magnification for representative images from tumor tissues. Morphometry based on stromal and tumor nest area measurements was performed by Olympus CellSens Dimensions Software package by manual annotation of measured areas, as previously described (Rizzardi et al, 2012). In the case of primary tumors, for one patient, 2 different TMA specimens were analyzed (A and B), retrieved from different regions of resected tumors. In the case of LN metastases, one TMA specimen was prepared from each LN sample. From all TMA blocks, 2 separate four-micron-thick sections (with a minimum of 100-micrometer distance in Z between them) were quantified using high resolution (20MP) 10× magnification images. Positive cells for immune markers CD45, CD3, CD8, and IDO, were identified by the presence of brown DAB precipitation around Hematoxylin-stained cell nuclei by a systematic quantitative method based on software-assisted, manual cell counting by two independent observers using the cell counter plugin of ImageJ software (Rueden et al, 2017). PVR expression was assessed semiquantitatively, where 0 = negative, 1 = low, 2 = moderate, 3 = strong, 4 = very strong expression scores were given for each specimen. Quantification of IDO expression was based on positive cell numbers in stroma and tumor nests in whole visual fields (10× magnification) of 2 separate sections of one TMA core. No DAB signs without the characteristic cellular shape or without the co-presence of nuclear staining were included in the calculations. Stromal and tumor nest total areas were measured using the area measurement tool in the Olympus CellSens Dimensions software package. Square micrometers (μm^2^) were converted to square millimeters (mm^2^) for calculation of cell density parameters in statistical analyses. Regions of apoptosis, necrosis and damage or disruptions in the sections were not included in the measurements. Results (cell numbers and areas) from separate sections of the same TMA punches were averaged before statistical assessment.

### Statistical Methods

We calculated differences using paired, 2-tailed t-test in stroma and tumor nest (tumor) area ratio or immune cell marker expressions (CD45, CD3, and CD8) in primary tumors and matched LN metastases, or according to NE-low vs NE-high subtypes in different anatomical localizations. Differences in stroma and tumor area ratio in primary tumors and LN metastases, according to NE subtypes, were also analyzed in selected cases with a one-way analysis of variance. Statistical analyses were performed on tumor samples with evaluable IHC data. Two-sided p-values less than 0.05 were considered statistically significant. Metric data were shown as mean, and corresponding SEM and graphs indicate the mean and corresponding 95% CI.

The association of immune checkpoints (PVR and IDO) and immune cell marker expression was evaluated using Spearman’s rank correlation. We assessed the linear correlation coefficient between the variables. The correlation coefficient (r) can vary between −1 to 1. No Correlation (0 < r ≤ 0.2), weak positive correlation (0.2 < r ≤ 0.5), moderate positive correlation (0.5 < r ≤ 0.8), strong positive correlation 0.8 < r < 1. All statistical analyses were implemented using the PASW Statistics 22.0 package (SPSS Inc., Chicago, IL, USA).

## RESULTS

In our study, we aimed to reveal the precise distribution pattern of immune cells *in situ* on SCLC tissue samples. For this, we performed IHC on serial sections of FFPE TMA samples, and demarcated the histological compartments of tumor stroma (stroma) and epithelial tumor nests (tumor) with consequent software-aided area measurement, followed by cell counting in every sample. First, we analyzed the histological distribution of immune cells in stroma vs. tumor nests in representative samples shown in **Fig 1**.

**Figure 1.**
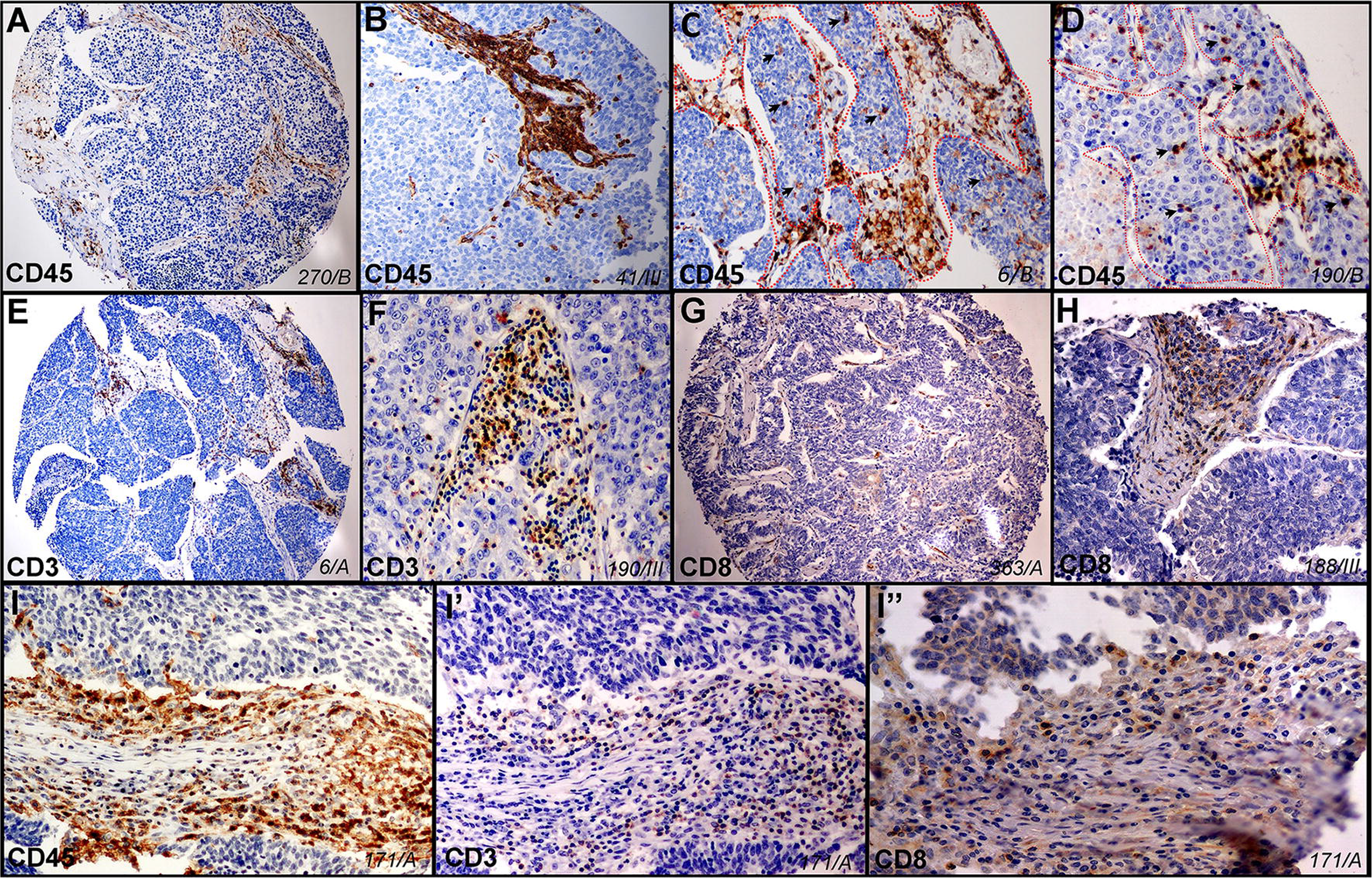
Histological localization of major immune cells in SCLC in representative tissue samples (ID of sample in italics). Qualitative *in situ* IHC data on the histological distribution of immune cells show high immune cell density in the stroma and a low number of labeled cells in tumor nests (**Fig 1A**, a magnified image in **1B**) stained with anti-CD45 antibody and hematoxylin. Infiltration of CD45+ immune cells in tumor nests can be low (**Fig 1A and B**) or moderate (**Fig 1C and D**), where dashed line signs the border of stroma and epithelial tumor nests (**Fig 1C and D)**and arrowheads show immune cells inside tumor nests (**Fig 1D**). Sections of whole TMA specimens stained with anti-CD3 and anti-CD8 antibodies show the presence of CD3+ T cells (**Fig 1E and F**) and CD8+ cytotoxic T cells (**Fig 1G and H**) in low (**Fig 1E and G**) and high (**Fig 1F and H**) magnification images in tumor stroma and sparsely in tumor nests. High magnification images of consecutive sections from the same TMA specimen and region of interest show CD45 (**Fig 1I**), CD3 (**Fig 1I’**) and CD8 (**Fig 1I’’**) labeling of tumor-infiltrating immune cells.

Pan-leukocyte marker CD45 labels every immunologically competent cell population from the myeloid and lymphoid lineage, but does not label erythrocytes. CD45 immunolabeling identifies a high number of immune cells in the stroma (**Fig 1A, B)**, but a limited number of cells in epithelial tumor nests **(Fig 1C and D**). Borders of fibrous stromal strands and tumor nests are shown with dashed lines, and immune cells inside tumor nests are indicated with arrowheads in **Fig 1C and D** on representative TMA sections. CD3 is the immunogen part of the T-cell receptor complex (TCR) that labels all mature T-cell populations of round cellular morphology (**Fig 1E and F**), whereas transmembrane glycoprotein CD8 represents the general marker for cytotoxic (effector) T cells (**Fig 1G and H**). Successive sections from the same primary tumor sample of SCLC patient show the expression of CD45 (**Fig 1I**), CD3 (**Fig 1 I’**) and CD8 (**Fig 1I’’**) on consecutively narrower cell populations (immune cells, T cells, cytotoxic T cells) in the same area of the TMA specimen.

Based on our *in situ* HE-stained sections, the stroma and tumor area ratio were similar in primary tumors (0.91±1.73) and LN (0.69±1.23) metastases (**S. Fig 1A**), and there were no statistically significant differences according to NE subtypes (**S. Fig 1B**).

### Immune cell distribution in primary tumors and lymph node (LN) metastases

Next, we compared the presence of immune cells according to anatomical localization. Immune cell marker expression according to primary tumors vs LN metastases is shown in **Fig 2**. We found that CD45+ (**Fig 2A and B**), CD3+ (**Fig 2E and F**), and CD8+ (**Fig 2I and J**) immune cell density was significantly higher in the stroma of LN metastases compared to primary tumors, but there was no significant difference in the case of tumor nests (tumor). Moreover, the stroma of primary tumors were significantly more infiltrated by major immune cells vs tumor nests, (CD45+, 912.0 ± 125.9 vs 65.83 ± 11.71 cell/mm^2^, respectively, p<0.01, **Fig 2C**; CD3+, 267.1 ± 45.98 vs 18.81 ± 4.125 cell/mm^2^, respectively, p<0.01, **Fig 2G**, and CD8+, 168.3 ± 30.54 vs 9.511 ± 2.409 cell/mm^2^, respectively, p<0.01, **Fig 2K**). Similarly, the stroma of LN metastases are significantly more infiltrated by major immune cells vs tumor nests (CD45+, 1993 ± 426.6 vs 107.9 ± 26.25 cell/mm^2^, respectively, p<0.01, **Fig 2D**; and CD3+, 585.4 ± 132.2 vs 39.50 ± 13.13 cell/mm^2^, respectively, p<0.01, **Fig 2H;** and CD8+, 356.0 ± 80.07 vs 21.62 ± 9.3 cell/mm^2^, respectively, p<0.01, **Fig 2L**).

**Figure 2.**
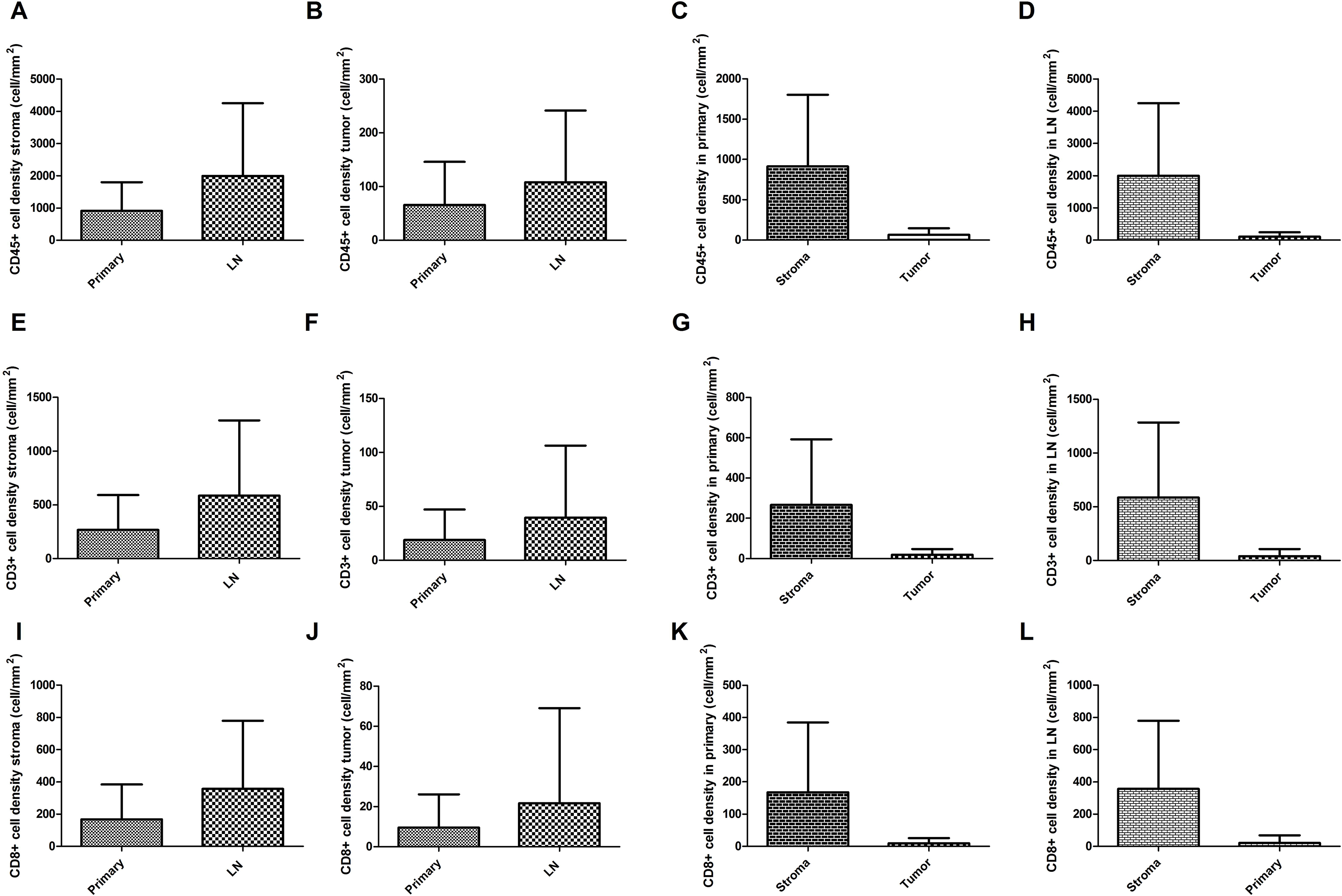
Immune cell distribution in primary SCLC tumors and matched lymph node (LN) metastases according to stroma and tumor nests. There were significantly higher cell density in the stroma of LN metastases compared to primary tumors, for CD45 (912.0 ± 125.9 vs 1993 ± 426.6, cell/mm^2^, p=0.0036, respectively, **Fig 2A**), CD3 (267.1 ± 45.98 vs 585.4 ± 132.2, cell/mm^2^, p=0.0076, respectively, **Fig 2E**), and CD8 (168.3 ± 30.54 vs 356.0 ± 80.07, cell/mm^2^, p=0.011, respectively, **Fig 2I**). Immune cell density showed no significant difference in tumor nests (tumor) in LN metastases compared to primary tumors, for CD45 (65.83 ± 11.71 vs 107.9 ± 26.25 cell/mm^2^, p= 0,0973, **Fig 2B**), CD3 (18.81 ± 4.125 vs 39.50 ± 13.13 cell/mm^2^, p= 0.0687, **Fig 2F**), and CD8 (9.511 ± 2.409 vs 21.62 ± 9.295 cell/mm^2^, respectively, p= 0.11, **Fig 2J**). Moreover, the stroma of primary tumors are significantly more infiltrated by major immune cells vs tumor nests, for CD45 (912.0 ± 125.9 vs 65.83 ± 11.71 cell/mm^2^, respectively, p< 0.0001, **Fig 2C**), CD3 (267.1 ± 45.98 vs 18.81 ± 4.125 cell/mm^2^, respectively, p < 0.0001, **Fig 2G**), and CD8 (168.3 ± 30.54 vs 9.511 ± 2.409 cell/mm^2^, respectively, p < 0.0001, **Fig 2K**). The stroma of LN metastases are significantly more infiltrated by major immune cells vs tumor nests, for CD45 (1993 ± 426.6 vs 107.9 ± 26.25 cell/mm^2^, respectively, p< 0.0001, **Fig 2D**), CD3 (585.4 ± 132.2 vs 39.50 ± 13.13 cell/mm^2^, respectively, p = 0.0002, **Fig 2H**), and CD8 (356.0 ± 80.07 vs 21.62 ± 9.295 cell/mm^2^, respectively, p < 0.0001, **Fig 2L**).

Next, we analyzed the relative distribution of major immune cells in stroma vs tumor nests, according to primary tumors and LN metastases (**S. Fig 2.**). There was no significant difference in CD45+/CD3+ cell ratio in stroma vs tumor nests when having pooled both primary and LN metastases (**S. Fig 2A**). Similarly, we found no significant difference in CD45+/CD3+ cell ratio in primary tumors vs LN metastases in the stroma (**S. Fig 2C**) or tumor nests (**S. Fig 2D**). In contrast, we found a significant difference in CD3+/CD8+ cell ratio in stroma vs tumor nests, when having pooled both primary and LN metastases (57.56 ± 2.469 vs 47.83 ± 4.410, p<0.01, respectively, **S. Fig 2B**). However, a separate analysis showed no statistically significant difference in CD3+/CD8+ ratio according to primary tumors and LN metastases in the stroma (**S. Fig 2E**) or tumor nests (**S. Fig 2F**).

### Immune cell distribution according to NE subtypes and tumor compartments

In primary tumors, we found a significantly increased density of CD45+ cells in NE-low compared to NE-high subtypes both in stroma (1359 ± 238.3 vs 660.7 ± 126.3, cell/mm^2^ respectively, p<0.01, **Fig 3A**) and tumor nests (tumor) (110.6 ± 24.95 vs 42.74 ± 10.30, cell/mm^2^ respectively, p<0.01, **Fig 3B**). Similarly, there were significantly increased density of CD3+ cells between NE-low and NE-high primary tumor subtypes both in stroma (418.9 ± 87.04 vs 181.7 ± 47.24, cell/mm^2^, respectively, p<0.01, **Fig 3E**) and tumor nests (35.50 ± 9.561 vs 10.19 ± 2.950, cell/mm^2^, respectively, p<0.01, **Fig 3F**). Correspondingly, there was a significantly increased density of CD8+ cells in NE-low primary tumors compared to NE-high subtypes both in stroma (301.2 ± 63.50 vs 93.56 ± 23.64, cell/mm^2^, respectively, p<0.01, **Fig 3I**) and tumor nests (21.81 ± 5.458 vs 3.161 ± 1.362, p<0.01, **Fig 3J**). Next, we analyzed LN metastases in terms of NE subtypes and immune cell distribution. Similarly to the primary tumors, we found a significantly increased density of CD45+ cells in NE-low LN metastases compared to NE-high subtypes in tumor nests (221.0 ± 78.06 vs 73.95 ± 20.86, cell/mm^2^, respectively, p= 0.0149, **Fig 3B**) but not in the stroma (**Fig 3A**).

**Figure 3.**
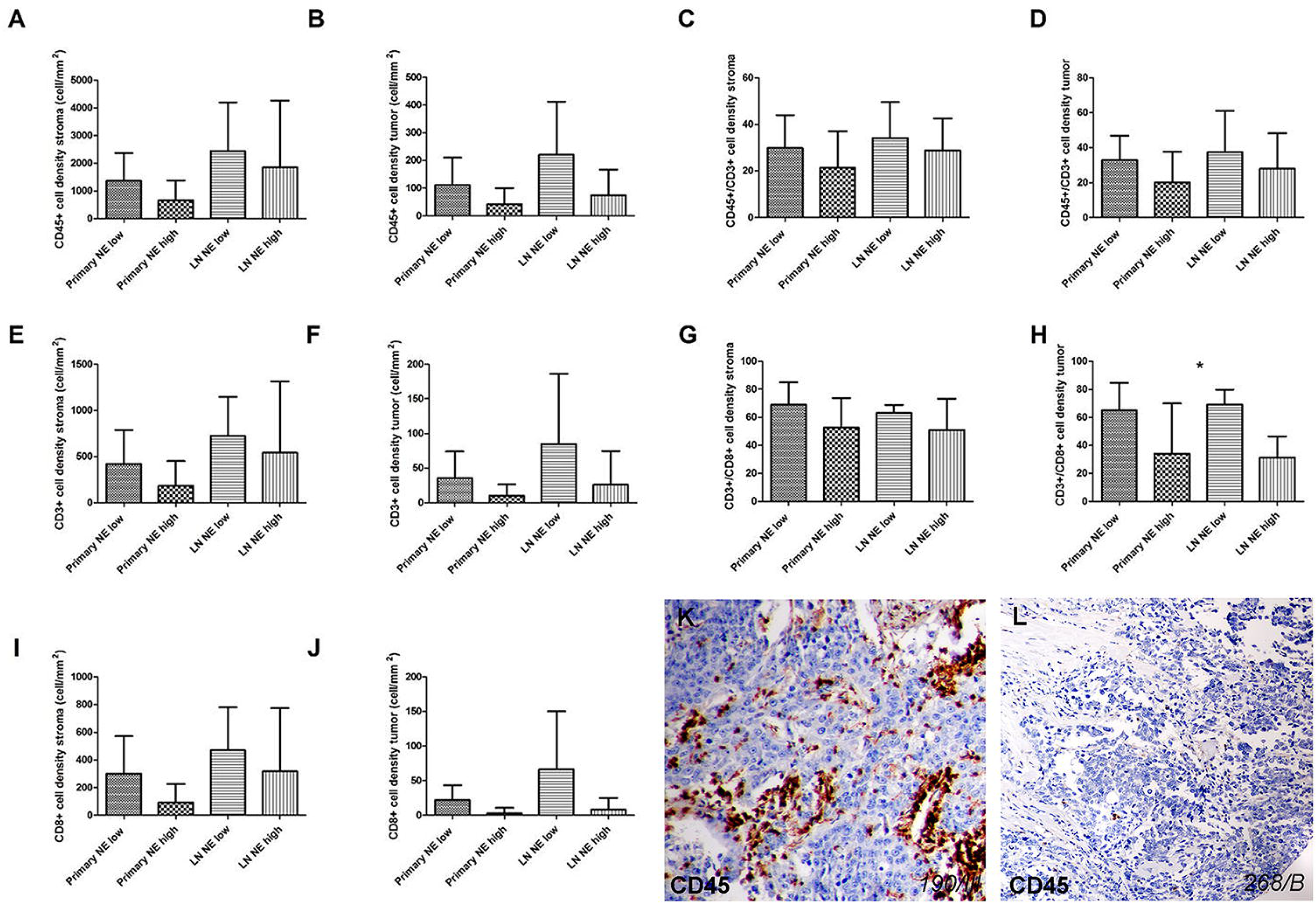
Immune cell distribution in primary SCLC tumors and matched lymph node (LN) metastases according to stroma and tumor nests based on neuroendocrine (NE) tumor subtypes. There were significantly increased CD45+ cell densities in NE-low primary tumor subtypes compared to NE-high ones including both stroma (1359 ± 238.3 vs 660.7 ± 126.3, cell/mm^2^ respectively, p=0.0064, **Fig 3A**) and tumor nests (tumor) (110.6 ± 24.95 vs 42.74 ± 10.30, cell/mm^2^ respectively, p= 0.0048, **Fig 3B**). We found a significantly increased density of CD45+ cells in NE-low LN metastases compared to NE-high subtypes in tumor nests (221.0 ± 78.06 vs 73.95 ± 20.86, cell/mm^2^, respectively, p= 0.0149, **Fig 3B**) but not in stroma (2436 ± 668.1 vs 1845 ± 527.7, cell/mm^2^, respectively, p= 0.5582, **Fig 3A**). There were significantly increased densities of CD3+ cells in NE-low primary tumor compared to NE-high subtypes both in stroma (418.9 ± 87.04 vs 181.7 ± 47.24, cell/mm^2^, respectively, p= 0.0117 **Fig 3E**) and tumor nests (35.50 ± 9.561 vs 10.19 ± 2.950, cell/mm^2^, respectively, p= 0.0026 **Fig 3F**). We found a significantly increased density of CD3+ cells in NE-low LN metastases compared to NE-high subtypes in tumor nests (84.67 ± 41.47 vs 25.95 ± 10.83, p= 0.05 **Fig 3F**) but not in stroma (721.9 ± 160.4 vs 539.9 ± 168.8, p= 0.5613 **Fig 3E**). There were significantly increased densities of CD8+ cells in NE-low primary tumor compared to NE-high subtypes both in stroma (301.2 ± 63.50 vs 93.56 ± 23.64, cell/mm^2^, respectively, p= 0.0006 **Fig 3I**) and tumor nests (21.81 ± 5.458 vs 3.161 ± 1.362, p < 0.0001 **Fig 3J**). We found a significantly increased density of CD8+ cells in NE-low LN metastases compared to NE-high subtypes in tumor nests (66.17 ± 34.30 vs 8.250 ± 3.765, cell/mm^2^, respectively, p= 0.0059 **Fig 3J**) but not in stroma (469.4 ± 117.6 vs 318.2 ± 99.36, cell/mm^2^, respectively, p= 0.4238 **Fig 3I**). According to NE-low and NE-high primary tumors the CD45+/CD3+ cell ratio was limited to 34.14 ± 5.845 % and 28.84 ± 3.156% (p=0.4069) in the stroma, and 32.85 ± 3.891% and 20.16 ± 4.017% (p=0.0376) in tumor nests (**Fig 3C and D**). There were no significant differences in CD45+/CD3+ cell ratio according to NE subtypes and tumor localization. According to NE-low and NE-high primary tumors the CD3+/CD8+ cell ratio was limited to 69.06 ± 3.758% and 52.58 ± 4.315% (p=0.0086) in the stroma and 64.85 ± 5.461% and 34.00 ± 10.38% (p=0.0131) in tumor nests, respectively (**Fig 3G and H**). According to NE-low and NE-high LN metastases the CD3+/CD8+ cell ratio was limited to 63.14 ± 2.143% and 50.89 ± 5.116%, p=0.1687 in the stroma and 69.00 ± 4.412% and 31.27 ± 4.545%, p< 0.0001 in tumor nests, respectively (**Fig 3G and H**). CD45 immunolabeling on a representative section of NE-low (**Fig 3K**) LN metastasis shows highly infiltrated stroma and tumor nests, whereas tumor-infiltrating immune cells are absent both in the stroma and the tumor nests on the sample of NE-high primary tumor (**Fig 3L**) (ID of the sample in italics).

Furthermore, we identified a significantly increased density of CD3+ cells in NE-low LN metastases compared to NE-high subtypes in tumor nests (84.67 ± 41.47 vs 25.95 ± 10.83, p= 0.05, **Fig 3F**) but not in the stroma (**Fig 3E**). Moreover, we found a significantly increased density of CD8+ cells in NE-low LN metastases compared to NE-high subtypes in tumor nests (66.17 ± 34.30 vs 8.25 ± 3.77, cell/mm^2^, respectively, p<0.01, **Fig 3J**) but not in the stroma (**Fig 3I**).

Next, we analyzed the relative immune cell distribution according to NE subtypes in stroma and tumor (**Fig 3 C, D, G, and H.**). The CD45+/CD3+ cell ratio was not significantly different in NE-low vs NE-high primary tumor stromata (**Fig 3C**). In contrast, the CD45+/CD3+ (32.85 ± 3.89% and 20.16 ± 4.02%, respectively, p=0.038, **Fig 3D**), and CD3+/CD8+ cell ratio (69.06 ± 3.76% and 52.58 ± 4.32%, respectively, p<0.01, p=0.038, **Fig 3H**) was significantly increased in primary tumor nests. Similarly to CD45+/CD3+ ratio in primary tumors, the CD3+/CD8+ cell ratio was not significantly increased in the stromal compartment of LN metastases (63.14 ± 2.14% and 50.89 ± 5.116%, p=0.17, **Fig 3G**). The CD3+/CD8+ cell ratio was significantly increased in NE-low tumor nests (compared to NE-high ones) (69 ± 4.41% and 31.27 ± 4.55%, p<0.01, respectively, **Fig 3H**) but there was no similar difference in term of stroma compartment (Fig 3G). **Fig 3K** shows a representative sample of NE-low SCLC subtype stained with CD45, where massive infiltration of stroma and a relatively high number of immune cells in tumor nests is characteristic. On the contrary, a typical ‘immune desert’ or infiltrate-excluded phenotype with scattered CD45+ cells both in stroma and tumor nests is shown in **Fig 3L** from a representative sample of NE-high SCLC tumor subtype.

### Immune checkpoint expression and NE subtypes

The expression pattern of emerging immune checkpoints Poliovirus receptor (PVR) and Indoleamine-2,3-dioxygenase 1 (IDO) in primary tumors and LN metastases according to NE-high vs NE-low tumors is shown in **Fig 4**. IHC shows that PVR is expressed by tumor cells, but not by stromal cells in both NE SCLC subtypes (**Fig 4A and B**). IDO is expressed by endothelial cells (**Fig 4D**) and stromal cells of various morphology (**Fig 4C**), just as by immune cells in tumor nests (**Fig 4E**) in both NE SCLC subtypes.

**Figure 4.**
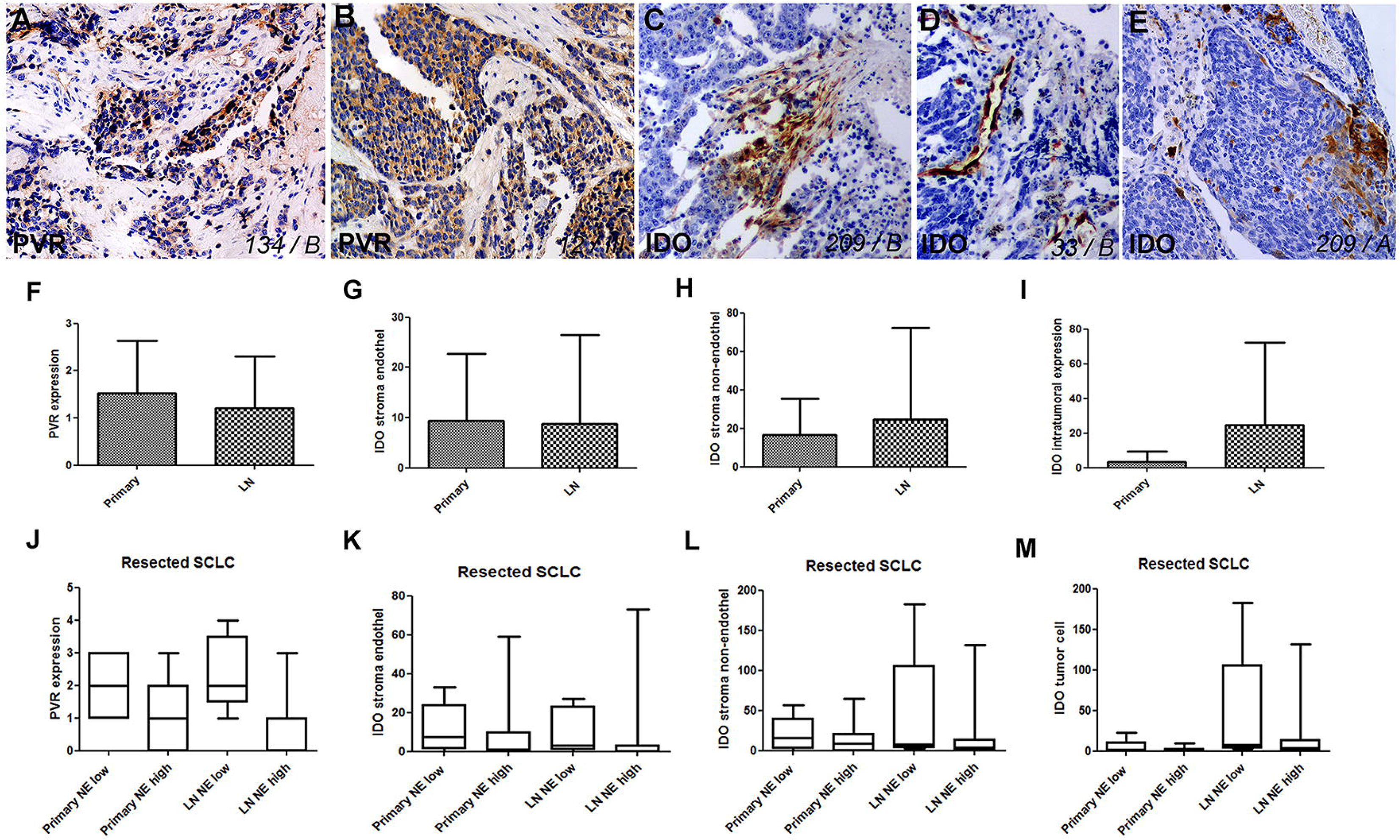
Distribution pattern of immune checkpoint Poliovirus receptor (PVR) and Indolamine-2,3-dioxygenase 1 (IDO) expression. PVR is expressed by tumor cells, but is not present in stromata including both NE SCLC subtypes (**Fig 4A and B**). IDO is expressed by endothelial cells (**Fig 4D**), stromal cells of various morphology (**Fig 4C**), and by immune cells in tumor nests (**Fig 4E**) in both NE SCLC subtypes. PVR expression showed no significant difference in primary tumor vs LN metastases (1.526 ± 0.1798 vs 1.217 ± 0.2263, p=0.2921, **Fig 4F**), but a significantly higher expression in NE-low vs NE-high was found both in primary tumors (2.200 ± 0.2225 vs 1. 087 ± 0.2170, p=0.0015, **Fig 4J**) and LN metastases (2.4000 ± 0.5099 vs 0.8889 ± 0.1962, p=0.0032, **Fig 4J**). There were no significant differences between primary tumors and LN metastases regarding the IDO expression of stroma endothelial (9.390 ± 2.092 vs 8.833 ± 3.615, p=0.8865, **Fig 4K**) and non-endothelial cells (16.63 ± 2.926 vs 24.79 ± 9.688, p=0. 3300, **Fig 4H**), whereas the intratumoral expression of IDO was higher in LN metastases compared to primary tumors (24.79 ± 9.688 vs 3.317 ± 0.9483, p=0.0055, **Fig 4I**). IDO stroma endothelium and non-endothelial cell expression showed no significant difference according to NE-low and NE-high tumor subtypes including primary tumors (11.50 ± 3.047 vs 8.296 ± 2.772, p=0.4747, **Fig 4K;** and 22.36 ± 5.517 vs 13.67 ± 3.335, p=0.1617, **Fig 4L**) and LN metastases (10.20 ± 5.398 vs 8.474 ± 4.400, p=0.8512 **Fig 4K;** and 45.60 ± 34.66 vs 19.32 ± 8.557, p=0.2802, **Fig 4L**). In contrast, the intratumoral expression of IDO was significantly higher in NE-low primary tumors (vs NE-high tumors) (6.786 ± 2.384 vs 1.519 ± 0.5130, p=0.0068, **Fig 4M**), but not in LN metastases (45.60 ± 34. 66 vs 19.32 ± 8.557, p=0.2802, **Fig 4M**).

PVR expression showed no significant difference in primary tumors vs LN metastases (**Fig 4F**). However, in NE-low subtype, a significantly higher expression was found compared to NE-high subtype both in primary tumors (2.2 ± 0.23 vs 1.1 ± 0.22, p<0.01, **Fig 4J**) and LN metastases (2.4 ± 0.51 vs 0.89 ± 0.19, p<0.01, **Fig 4J**).

There was no significant difference in IDO expression of stroma endothelial (**Fig 4G**) and non-endothelial cells (**Fig 4H**) in primary tumors and LN metastases. In contrast, the intratumoral expression of IDO was higher by orders of magnitude in LN metastases compared to primary tumors (24.79 ± 9.69 vs 3.32 ± 0.95, p<0.01, **Fig 4I**). Next, we assessed IDO expression on different cell types and in different tumor compartments regarding the NE phenotype. IDO stroma endothelium and non-endothelial cell expression showed no significant difference between NE-low and NE-high tumor subtypes neither in primary tumors (**Fig 4K and 4L**) nor in LN metastases (**Fig 4K and 4L**). On the contrary, intratumoral expression of IDO was significantly higher in NE-low (vs NE-high) primary tumors, (6.8 ± 2.39 vs 1.52 ± 0.51, p<0.01, **Fig 4M**), but not in LN metastases (**Fig 4M**).

Next, we investigated the associations between the expression of immune checkpoints and immune cell infiltration, where Spearman’s correlation coefficient was calculated (Table 1). We found a significant moderate positive correlation between IDO stroma endothelium, stroma non-endothelial cell and immune cell density in stroma including CD45+ cells (r= 0.7, r= 0.682, respectively, p<0.01, **Fig 5 A and B**), and CD8+ T cells (r= 0.783, r= 0.727, respectively, p<0.01, **Fig 5 D and E**) in primary tumors. Furthermore, there was a statistically significant strong positive correlation between primary tumor IDO intratumoral expression and immune cell density in tumor nests, including CD45+ cells (r= 0.755, p<0.01, **Fig 5C**), and CD8+ T cells (r= 0.763, p<0.01, **Fig 5F**).

**Figure 5.**
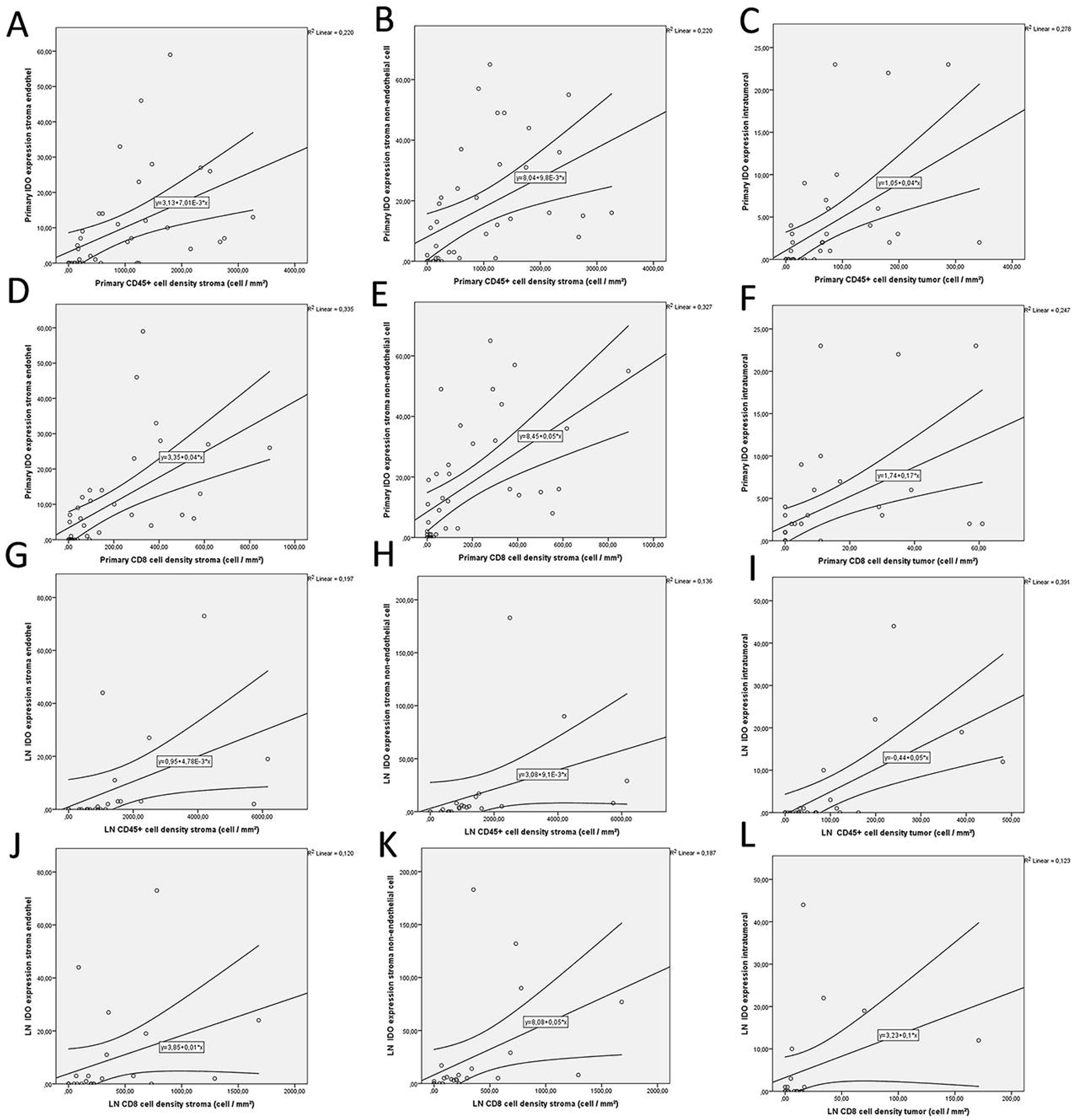
Plot diagrams of significant moderate-to-strong correlations between immune checkpoints and immune cell infiltration. There was a significant moderate positive correlation between primary tumor IDO stroma endothelium, stroma non-endothelial cell and immune cell density in stroma including CD45+ cells (r= 0.7, r= 0.682, respectively, p<0.01, **Fig 5 A and B**), and CD8+ T cells (r= 0.783, r= 0.727, respectively, p<0.01, **Fig 5 D and E**). Similarly, there was a statistically significant strong positive correlation between primary tumor IDO intratumoral expression and immune cell density in tumor nests, including CD45+ cells (r= 0.755, p<0.01, **Fig 5C**), and CD8+ T cells (r= 0.763, p<0.01, **Fig 5F**). A statistically significant strong positive correlation was present between IDO stroma endothelium, stroma non-endothelial cell, and CD45+ cell density in the stroma of LN metastases (r= 0.806, r= 0.835, respectively, p<0.01, **Fig 5 G and H**). There was a significant moderate positive correlation between IDO stroma endothelium, stroma non-endothelial cell, and CD8+ T cell density in the stromata of LN metastases (r= 0.542, P<0.05, r= 0.748, respectively, p<0.01, **Fig 5 J and K**). Moreover, there was a statistically significant strong positive correlation between primary tumor IDO intratumoral expression and immune cell density in tumor nests, including CD45+ cells (r= 0.689, p<0.01, **Fig 5I**), and CD8+ T cells (r= 0.581, p<0.01, **Fig 5L**).

**Table 1.** Correlation of immune checkpoints expression and immune cell infiltration. r: Correlation Coefficient, p: probability value, N: number of patients, Primary: primary tumor, LN: Lymph node metastasis, Stroma: Tumor stroma area, Tumor: Tumor nest including intratumoral area

|  |  |  | Primary tumor cell density (cell / mm<sup>2</sup>) |  |  |  |  |  | Lymph node metastasis cell density (cell / mm<sup>2</sup>) |  |  |  |  |  |
| --- | --- | --- | --- | --- | --- | --- | --- | --- | --- | --- | --- | --- | --- | --- |
|  |  |  | CD45+ |  | CD3+ |  | CD8+ |  | CD45+ |  | CD3+ |  | CD8+ |  |
|  |  |  | stroma | tumor | stroma | tumor | stroma | tumor | stroma | tumor | stroma | tumor | stroma | tumor |
| Primary | PVR expression | r | ,468** | ,550** | ,448** | ,435** | ,481** | ,552** | 0,25 | 0,12 | 0,34 | 0,20 | 0,30 | 0,21 |
|  |  | p | 0,00 | 0,00 | 0,00 | 0,01 | 0,00 | 0,00 | 0,32 | 0,64 | 0,16 | 0,42 | 0,22 | 0,41 |
|  |  | N | 38,00 | 38,00 | 38,00 | 38,00 | 38,00 | 38,00 | 18,00 | 18,00 | 19,00 | 18,00 | 19,00 | 18,00 |
|  | IDO stroma endothel | r | ,700** | ,812** | ,771** | ,845** | ,783** | ,737** | -0,25 | -0,04 | -0,07 | -0,01 | -0,03 | 0,04 |
|  |  | p | 0,00 | 0,00 | 0,00 | 0,00 | 0,00 | 0,00 | 0,31 | 0,86 | 0,75 | 0,96 | 0,90 | 0,88 |
|  |  | N | 40,00 | 39,00 | 40,00 | 39,00 | 40,00 | 39,00 | 19,00 | 18,00 | 20,00 | 18,00 | 20,00 | 18,00 |
|  | IDO stroma non-endothelial cell | r | ,682** | ,794** | ,747** | ,808** | ,727** | ,710** | -0,22 | -0,06 | -0,03 | 0,01 | 0,02 | 0,10 |
|  |  | p | 0,00 | 0,00 | 0,00 | 0,00 | 0,00 | 0,00 | 0,37 | 0,82 | 0,90 | 0,95 | 0,92 | 0,69 |
|  |  | N | 40,00 | 39,00 | 40,00 | 39,00 | 40,00 | 39,00 | 19,00 | 18,00 | 20,00 | 18,00 | 20,00 | 18,00 |
|  | IDO tumor | r | ,676** | ,755** | ,747** | ,795** | ,737** | ,763** | -0,04 | 0,24 | 0,12 | 0,33 | 0,16 | 0,29 |
|  |  | p | 0,00 | 0,00 | 0,00 | 0,00 | 0,00 | 0,00 | 0,86 | 0,34 | 0,61 | 0,18 | 0,49 | 0,25 |
|  |  | N | 40,00 | 39,00 | 40,00 | 39,00 | 40,00 | 39,00 | 19,00 | 18,00 | 20,00 | 18,00 | 20,00 | 18,00 |
| LN | PVR expression | r | -0,10 | -0,03 | -0,12 | -0,10 | -0,12 | -0,25 | ,658** | ,577** | ,419* | ,608** | 0,38 | ,574** |
|  |  | p | 0,62 | 0,90 | 0,57 | 0,64 | 0,56 | 0,23 | 0,00 | 0,00 | 0,04 | 0,00 | 0,06 | 0,00 |
|  |  | N | 25,00 | 24,00 | 25,00 | 24,00 | 25,00 | 24,00 | 23,00 | 24,00 | 24,00 | 24,00 | 25,00 | 24,00 |
|  | IDO stroma endothel | r | -0,39 | -0,08 | -0,20 | 0,09 | -0,17 | -0,07 | ,806** | ,641** | ,450* | ,515* | ,542** | ,523* |
|  |  | p | 0,06 | 0,72 | 0,34 | 0,67 | 0,42 | 0,76 | 0,00 | 0,00 | 0,03 | 0,01 | 0,01 | 0,01 |
|  |  | N | 24,00 | 23,00 | 24,00 | 23,00 | 24,00 | 23,00 | 22,00 | 23,00 | 23,00 | 23,00 | 24,00 | 23,00 |
|  | IDO stroma non-endothelial cell | r | -0,22 | -0,06 | -0,09 | 0,10 | -0,02 | 0,17 | ,835** | ,779** | ,730** | ,665** | ,748** | ,679** |
|  |  | p | 0,30 | 0,77 | 0,69 | 0,63 | 0,91 | 0,43 | 0,00 | 0,00 | 0,00 | 0,00 | 0,00 | 0,00 |
|  |  | N | 24,00 | 23,00 | 24,00 | 23,00 | 24,00 | 23,00 | 22,00 | 23,00 | 23,00 | 23,00 | 24,00 | 23,00 |
|  | IDO tumor | r | -0,22 | 0,01 | -0,06 | 0,19 | -0,05 | -0,05 | ,661** | ,689** | 0,30 | ,558** | 0,40 | ,581** |
|  |  | p | 0,31 | 0,98 | 0,78 | 0,39 | 0,81 | 0,81 | 0,00 | 0,00 | 0,17 | 0,01 | 0,05 | 0,00 |
|  |  | N | 24,00 | 23,00 | 24,00 | 23,00 | 24,00 | 23,00 | 22,00 | 23,00 | 23,00 | 23,00 | 24,00 | 23,00 |

\*Correlation is significant at the 0.05 level (2-tailed Spearman test).
\*\*Correlation is significant at the 0.01 level (2-tailed Spearman test).

In terms of LN metastases, we found a statistically significant strong positive correlation between IDO stroma endothelium, stroma non-endothelial cell, and CD45+ cell density in stroma (r= 0.806, r= 0.835, respectively, p<0.01, **Fig 5 G and H**). Additionally, we found a significant moderate positive correlation between IDO stroma endothelium, stroma non-endothelial cell and CD8+ T-cell density in stroma (r= 0.542, P<0.05, r= 0.748, respectively, p<0.01, **Fig 5 J and K**). Moreover, there was a statistically significant strong positive correlation between IDO intratumoral expression and immune cell density in tumor nests including CD45+ cells (r= 0.689, p<0.01, **Fig 5I**), and CD8+ T cells (r= 0.581, p<0.01, **Fig 5L**).

Additionally, we found a statistically significant moderate positive correlation between primary tumor PVR expression and immune cell density both in tumor nests and matched LN metastases, including CD45+ cells (**S. Fig 3A and C**), and CD8+ T cells (**Fig 3B and D**). **Supplemental Figure 3.** shows plot diagrams of statistically significant moderate positive correlations, including both primary tumor and LN metastases, tumor PVR expression, and immune cell density.

## DISCUSSION

The standard of care therapy for extensive-stage SCLC now includes immunotherapy in the front-line setting. The addition of atezolizumab or durvalumab to chemotherapy has changed practice recently, and is associated with a moderate significantly longer PFS and OS than chemotherapy on its own (Horn et al, 2018; Pas-Ares et al, 2019). However, as of yet, no predictive biomarkers have been identified, and the PFS curves seem to overlap during the initial 8 months, showing that most patients do not benefit from immunotherapy. Additionally, there were increased benefits for selected patients that might respond to immunotherapy.

Recent advancements in transcriptomics studies highlight the potential of a distinct microenvironment in SCLC NE subtypes. Understanding the immunology of NE subtypes might affect the clinical outcome and help lay the framework for immunotherapy administration in this devastating cancer (Calbo et al, 2011; George et al, 2015; McColl et al, 2017). However, to date, studies have been performed exclusively on NSCLC samples. Therefore, our study aims to fill this gap of knowledge with a detailed IHC analysis of immune cell populations on human SCLC tissue samples. We aim to provide an in-depth inter-tumor heterogeneity array of IHC staining on primary tumors versus matched LN metastases on immune cell infiltration and immune activation of stroma and epithelial tumor nests in NE-low and NE-high tumor phenotypes. Importantly, to our knowledge, this is the first human study delivering *in situ* proteomics data on immune cell populations in LN metastases of SCLC patients.

Our main findings from this study interpret the proteomic profile of the tumor microenvironment to further highlight the relevance of NE-low vs. NE-high tumor subtypes in the clinical setting. It is also important to note that the presence of lymphatics-associated genes might influence any transcriptomic study performed on LN metastases. Therefore, our *in situ* proteomic analysis might overcome the limitations above.

Others showed that the extent of immunological infiltration in tumor tissue and the expression of immune checkpoints proved to be a reliable marker for response to anti-PD immunotherapy and long term survival in SCLC (Muppa et al., 2019; Bonanno et al., 2018); NSCLC (Al-Shibli et al, 2008), and other malignancies, like breast cancer (Bates et al, 2006), melanoma (Clemente et al, 1996), colorectal carcinoma (Galon et al, 2006), and prostatic cancer (Vesalainen et al, 1994) as well. Another group indicated on NSCLC TMA samples that a high number of stromal CD4+ and epithelial and stromal CD8+ cells were independent positive prognostic markers, and CD8+ Tumor-Infiltrating Lymphocytes (TILs) can stratify immunotherapy-treated patients of different clinical outcome (Lin Z et al., 2019).

Furthermore, a low level of CD8+ lymphocyte infiltration in tumor stroma was positively correlated with an augmented incidence of angiolymphatic tumor invasion (Al-Shibli et al., 2008). Additionally, fluorescence-activated cell sorting (FACS) analysis demonstrated that a special subset of CD103+/CD8+ T cells displayed increased activation-induced apoptosis in NSCLC and mediated cytolytic activity toward autologous tumor cells inhibiting PD-1 and PD-L1 interaction (Djenidi et al., 2015). However, another study in NSCLC patients suggests that CD8+ T cells in tumor islets can show immunological tolerance against tumor cells with increased expression of PD-1 and impaired immune function, including reduced production of cytokines and proliferation impairment (Zhang et al., 2010). Based on IHC data, SCLC patients with brain metastases and an increased number of TILs have significantly more prolonged survival (Berghoff et al., 2016).

First, we revealed that immune cell infiltration both in primary tumors and LN metastases are predominant in loosely arranged stromal bands, but not in tumor nests. Even in selected, relatively highly infiltrated tumors, only about 7% in primary and 5% in LN metastasis of CD45+ cells and 5% and 6%, respectively of CD8+ T cells are localized in the close microenvironment of tumor cells (**Fig 2C, D, G, H, K, and L**). Furthermore, we established that the stroma of LN metastases had significantly higher immune cell density compared to primary tumors; however, this difference was not significant in tumor nests (**Fig 2A, B, E, F, I, and J**).

Others found that decreased tumor/stroma ratio proved to be an independent negative prognostic factor in colon (Mesker et al, 2009; Huijbers et al, 2013), breast (Roeke et al, 2017), oesophageal (Wang et al, 2012), and lung cancer (Zhang et al, 2015; Wang et al, 2013). In SCLC, no data on tumor/stroma ratio was reported, and in vitro morphological characteristics and variance in cellularity were shown to be associated with NE phenotypes (Gazdar, 2018; Saito et al., 2018). In our study, we found no significant difference in terms of tumor/stroma ratio according to NE pattern neither in primary tumors nor in LN metastases (**S Fig 1B**). Furthermore, despite the significantly higher stroma immune cell density, LN metastases are not significantly more stromatous compared to primary tumors (**S Fig 1A**).

Our analyses demonstrate that both stromal and intratumoral CD45+/CD3+ cell ratio is limited to 27% when pooling both primary tumors and LN metastases (**S Fig 2A**). This means that TILs make up only about one quarter of all immune cells regardless of their anatomic (macroscopic) localization (primary tumor vs LN). Consequently, a significant fraction of CD45+ cells belongs to populations of macrophages, dendritic cells, neutrophils, or other non-specific immune cells in SCLC. On the other hand, our results showed that around half of the lymphocytes (and ~13% of CD45+ cells) are CD8+ cytotoxic T cells (**S Fig 2B, E, and F**) that correspond with FACS data (AACR 2019). In contrast to CD45+/CD3+ cell ratios, we found a significant difference in CD3+/CD8+ cell ratio in stroma vs tumor when pooled both primary and LN metastases, meaning tumor stroma has a significantly higher ratio of effector T cells compared to tumor nests (p=0.038, **S Fig 2B**).

Nevertheless, the latest SCLC clinical studies showed no difference in intraepithelial CD3+ infiltration between patients with shorter vs. longer survival. However, significantly higher CD3+ infiltration in the stroma and at the cell-cell interface was present among patients with more prolonged survival (Muppa et al, 2019). Immune response evasion by tumors may play a crucial role in the aggressiveness of SCLC, and can be due to the insufficient number of T cells or T cells that are inhibited in the tumor microenvironment (Antonia et al., WCLC 2019).

Our group on the same TMA sets clustered NE-high and NE-low SCLC subsets (Lohinai et al, 2020) based on the top RNA genes associated with NE differentiation (Gazdar 2018; Zhang et al., 2018). Furthermore, the latest preclinical studies suggest that, compared to the NE-high, the NE-low subtype is more likely to respond to immunotherapy due to its ‘immune oasis’ phenotype, emphasizing the necessity and importance of molecular and *in situ* immunological characterization before the assessment of therapies to this type of recalcitrant cancer (Gazdar 2018; Zhang et al., 2018; Ito et al., 2016).

Next, we compared *in situ* the quantitative and qualitative extent of the immunological microenvironment of SCLC tumors according to NE-low and NE-high subtypes. In line with previously published *in vitro* data, our results confirm that NE-low tumors are significantly more infiltrated by immune cells, primarily by CD8+ effector T cells (Zhang et al., 2018). Interestingly, in our study, the CD45+/CD3+ cell ratio was not significantly different in NE-low relative to NE-high tumors, suggesting that the T-cell population is not predominant neither in stroma nor in tumor nests in NE-low tumors. In contrast, a substantially higher percentage of CD8-expressing lymphocytes are present both in NE-low primary tumors and LN metastases (vs NE-high), and the difference is even more considerable in tumor nests.

Next, in order to identify targets and further understand the immune microenvironment, we analyzed the immune checkpoint expressions. PVR (CD155) has been reported to mediate T cell activation via CD226, or impede T-lymphocytes by binding to TIGIT. PVR overexpression is associated with poor prognosis in melanoma, colorectal, lung, and pancreatic cancers (Masson et al., 2001; Nakai et al., 2010; Bevelacqua et al., 2012; Nishiwada et al., 2015). Our data show that strong PVR expression was significantly more frequent in NE-low vs high tumors, including both primary tumors and LN metastases. Although PVR overexpression was correlated with poor prognosis in multiple studies (Nakai et al., 2010; Nishiwada et al., 2015), we found a moderate positive correlation between PVR expression and immune cell density in SCLC tumor nests including CD45+ (r= 0.55, p<0.01), and CD8+ T cells (r= 0.552, p<0.01). Of note, the latter finding was statistically significant in primary tumors and also in LN metastases.

Another checkpoint, IDO, originally belonged to anti-cancer molecules based on its anti-pathogenic function (Schmidt et al., 2014). Subsequent studies, however, identified tissue macrophages producing high levels of IDO upon IFN-γ stimulation inhibiting effector T-cell proliferation (Munn et al., 1998). IDO expression was reported in lung cancer cell lines (Karanikas et al., 2007), and *in situ* in 42-43% of NSCLC samples (Schalper et al., 2017; Volaric et al., 2018). We found IDO expression on stromal cells of various morphology. Of note, the presence of IDO in stroma endothelial cells is a novel finding in SCLC. Previous studies showed that the endothelial expression of IDO in metastatic kidney cancer promotes response to immunotherapy and is associated with better progression-free survival (Seeber et al., 2018). In line with other NSCLC studies, we observed scattered IDO immunolabeling only on tumor nest immune cells but not on tumor cells (Volaric et al., 2018). The role of IDO was demonstrated in other respiratory conditions as well, like pneumonia, where inflammatory macrophages were identified as a primary source of the molecule (Virok et al., 2019). IDO expression was not different in stromal cellular elements or endothelium according to NE subtypes. However, IDO expression in tumor nests showed significantly higher levels in NE-low tumors. Consequently, establishing an immunosuppressive microenvironment for TILs, that might explain why NE-low tumors do not unequivocally have better prognosis despite their ‘immune oasis’ phenotype. Stroma IDO expression might be associated with many types of inhibitory cells in the immunosuppressive tumor microenvironment, like cancer-associated fibroblasts, myeloid-derived suppressor cells (MDSC), or tumor-associated macrophages, which requires further confirmation. Interestingly, intratumoral expression of IDO showed a conspicuous discrepancy in LN metastases where IDO positive cells were much more abundant, than in primary tumors (24,79 ± 9,688 vs 3,317 ± 0,9483, p=0,0055, **Fig 4I**). Our findings show that LN metastases are significantly more infiltrated by immune cells (vs primary tumors). This might result in clinically indifferent molecular behavior and aggressiveness of LN metastases due to their distinct immunological microenvironment and immune checkpoint expression patterns.

Next, we analyzed the correlation of IDO and *in situ* immune cell infiltration. We hypothesize that a statistically significant strong positive correlation exists between intratumoral expression of IDO and immune cell density in tumor nests including CD45+ cells (**Fig 5C**), and CD8+ T cells (**Fig 5F**). Our data suggests that IDO overexpression is an escape mechanism of tumor cells making immune cells and lymphocytes entering tumor nests anergic and unable to launch an immune response against them.

This study has limitations. First of all, this is a retrospective cross-sectional study with limited clinicopathological data available. The patient population is unique in terms of the resected sample size and corresponding LN availability, however, it is small even in the light of the fact that matched tumor samples are usually not available in the case of SCLC. Prognostic data is limited, and our study may be influenced by the intraoperative techniques and differences in administration of oncotherapy over a long period. Protein expression was examined by antibodies; consequently, results might slightly vary according to the aspecific antibody used.

## Conclusion

To our knowledge, this is the first human study that demonstrates that SCLC stroma is more infiltrated by immune cells compared to tumor nests. Additionally, NE-low tumors are more infiltrated by immune cells compared to NE-high tumors. Therefore, our results suggest that SCLC is classified into two distinct NE subtypes that may alter treatment outcomes. Accordingly, we hypothesize that NE-low tumors have a microenvironment potentially associated with increased benefit from immune-checkpoint inhibitor therapy. Consequently, NE subtypes should be defined in future clinical trials to select patients that most likely benefit from immunotherapy administration. Moreover, PVR and IDO are potential new targets in SCLC with increased expression in NE-low subtype, providing critical insight for further prospective studies on SCLC immunotherapies.

## Acknowledgments

The authors thank the patients and the clinical teams involved. We thank Leslie Rozeboom for her assistance in creating TMA blocks.

---

**Supplementary Figure 1. Tumor nest and stroma area ratio in primary SCLC tumors and matched lymph node (LN) metastases, according to neuroendocrine (NE) tumor subtypes.**

There were no significant differences in stroma and tumor nest (tumor) area ratio in primary tumors and matched LN (0.91 ± 1.73 vs 0.69 ± 1.23, respectively, p=0.57; paired, 2-tailed t-test) metastases (**S. Fig 1A**). No significant differences were present in stroma and tumor area ratio in primary tumors and LN metastases according to NE subtypes (one-way analysis of variance, p=0.6; Primary NE-low vs high, 0.5147 ± 0.094 vs 1.115 ± 0.3757, respectively, p=0.17; LN NE-low vs high, 0.6915 ± 0.25 vs 0.6878 ± 0.31, respectively, p= 0.6878 ± 0.30 **S. Fig 1B**).

**Supplementary Figure 2. Relative distribution of immune cells according to primary SCLC tumors and matched LN metastases.**

There was no significant difference in CD45+/CD3+ cell ratio between stroma tumor when pooling both primary and LN metastases (26.72 ± 1.79 vs 27.45 ± 2.55, respectively, p=0.8, **S. Fig 2A**). In contrast, we found a significant difference in CD3+/CD8+ cell ratio between stroma and tumor when pooling both primary and LN metastases (57.56 ± 2.469 vs 47.83 ± 4.410, p=0.0398, respectively, **S. Fig 2B**). There was no significant difference in CD45+/CD3+ cell ratio in stroma according to primary tumors and LN metastases (24.72 ± 2.285, vs 30.27 ± 2.769, p=0.14, respectively, **S. Fig 2C**). We found no significant difference in CD45+/CD3+ cell ratio in tumor according to primary tumors and LN metastases (25.31 ± 3.033 vs 30.43 ± 4.420, p=0.32, respectively, **S. Fig 2D**). There was no significant difference in CD3+/CD8+ ratio in stroma according to primary tumors and LN metastases (59.64 ± 3.178, vs 54.19 ± 3.905, p=0.28, respectively, **S. Fig 2E**). No significant difference was identified in CD3+/CD8+ ratio in tumor according to primary tumors and LN metastases (50.04 ± 6.429 vs 44.59 ± 5.551, p=0.55, respectively, **S. Fig 2F**).

**Supplementary Figure 3. Plot diagrams of significant moderate correlations between the poliovirus receptor (PVR) and immune cell infiltration.**

There was a statistically significant moderate positive correlation between primary tumor PVR expression and immune cell density in tumor nests including CD45+ cells (r= 0.55, p<0.01), and CD8+ T cells (r= 0.552, p<0.01). Furthermore, in terms of LN metastases, a similarly moderate significant positive correlation was found between primary tumor PVR expression and immune cell density in tumor nests, including CD45+ cells (r= 0.577, p<0.01), and CD8+ T cells (r= 0.574, p<0.01).

