## Supplementary figures and images for "Small Cell Lung Cancer Neuroendocrine Subtypes are Associated with Different Immune Microenvironment and Checkpoint Molecule Distribution"

### Supplemental Figure 1

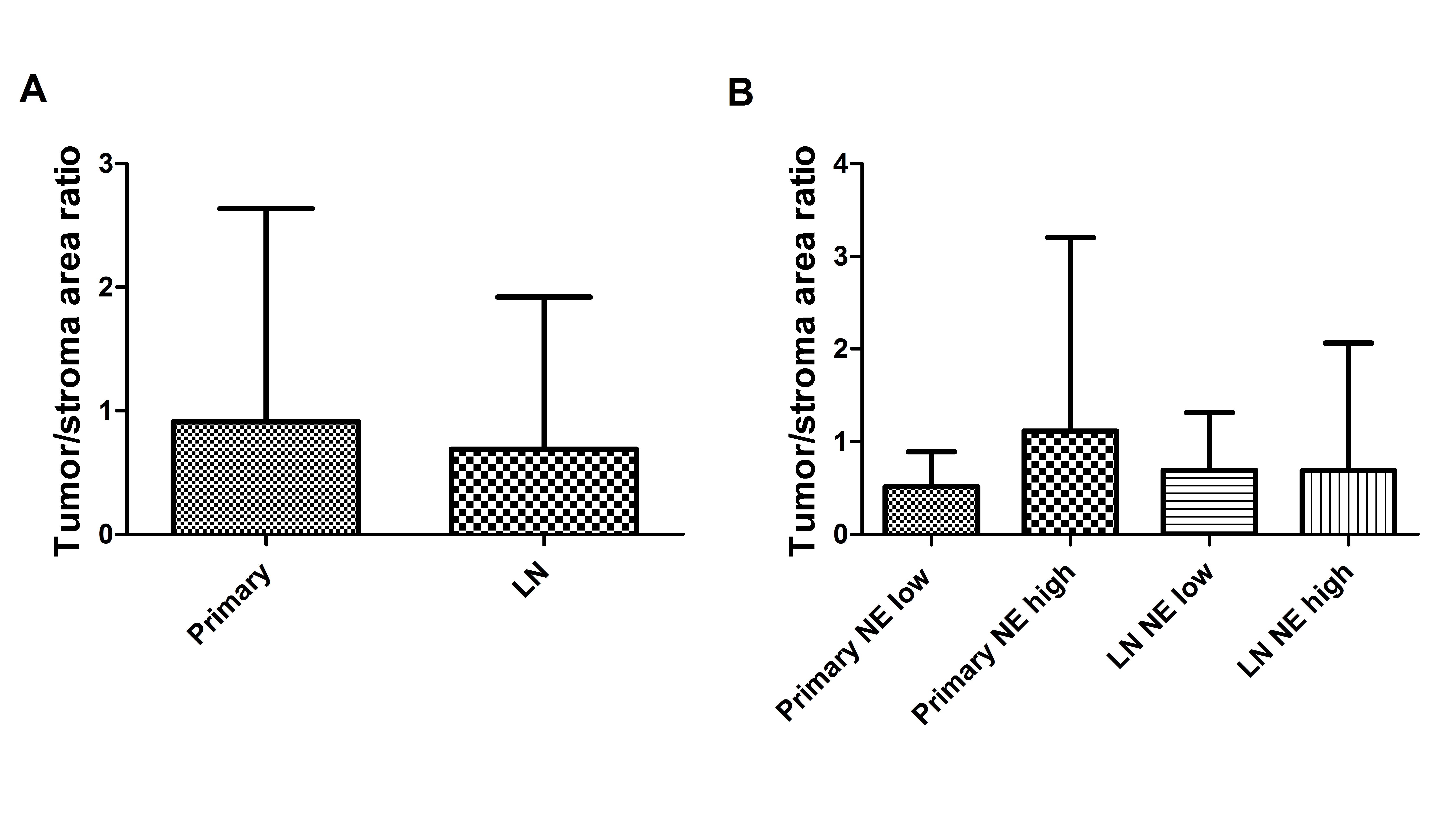

### Supplemental Figure 2

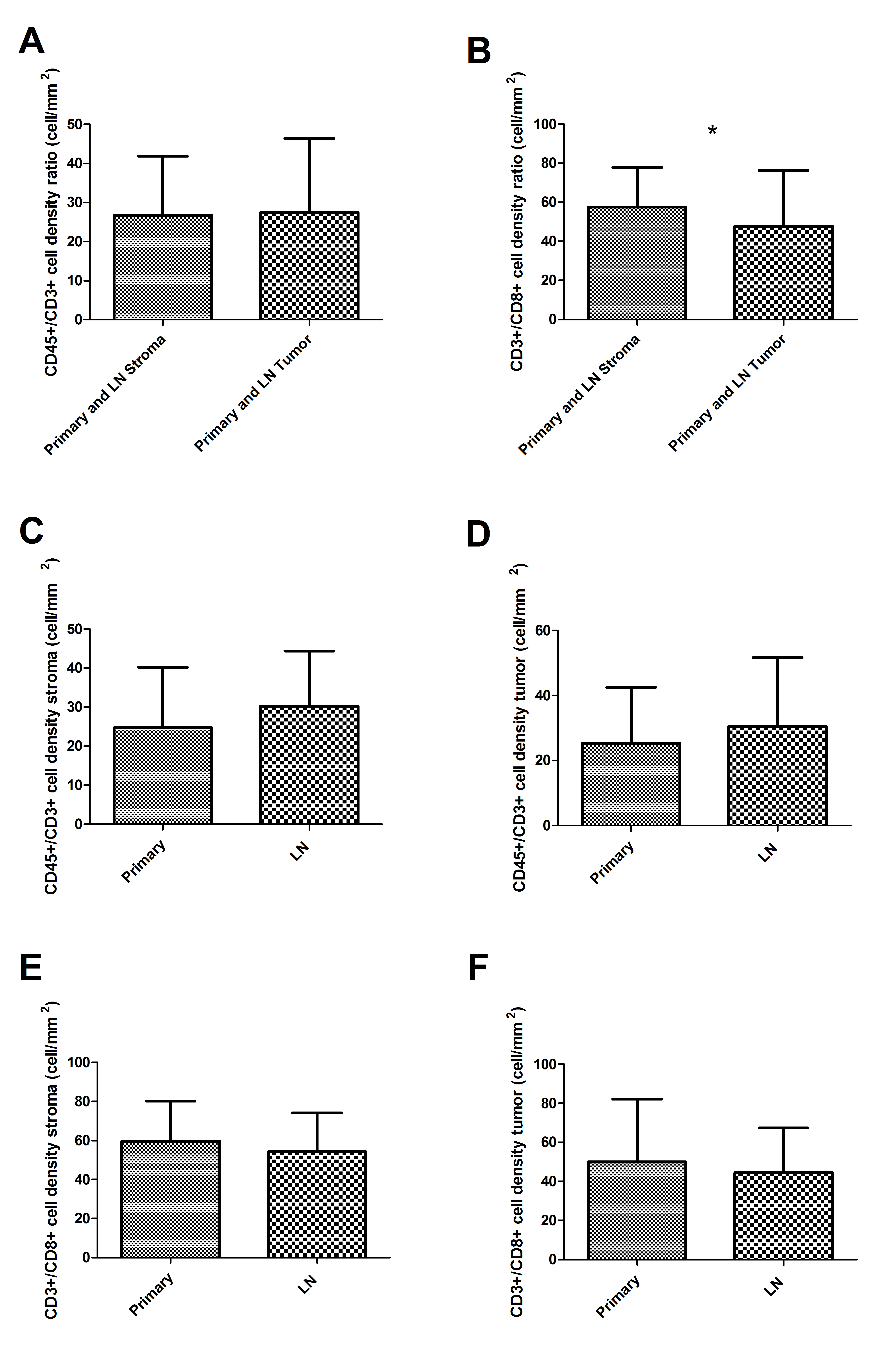

### Supplemental Figure 3

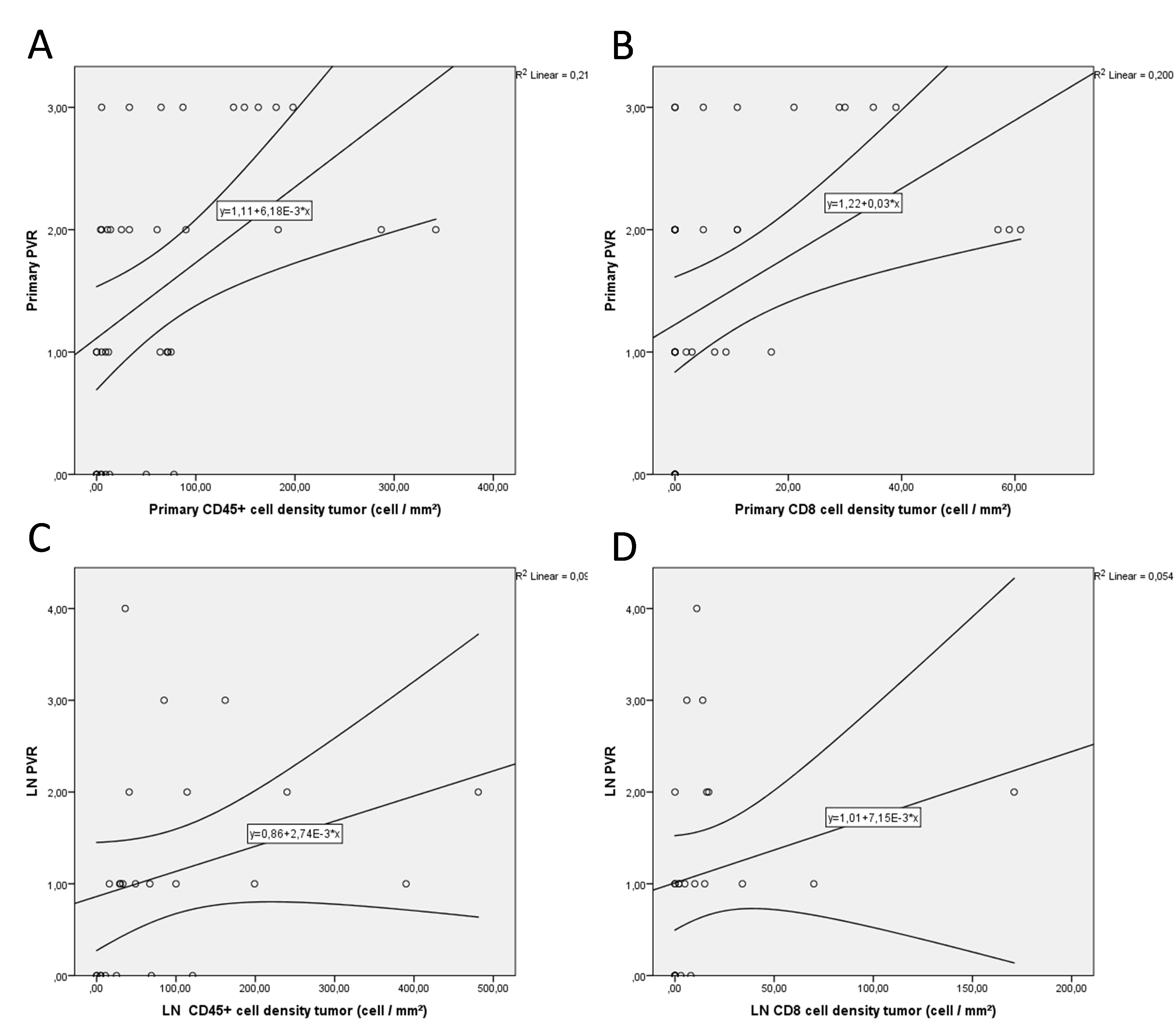
